# Active antithrombin glycoforms are selectively physiosorbed on plasma extracellular vesicles

**DOI:** 10.1101/2021.07.16.452649

**Authors:** Annalisa Radeghieri, Silvia Alacqua, Andrea Zendrini, Vanessa Previcini, Francesca Todaro, Giuliana Martini, Doris Ricotta, Paolo Bergese

**Author notes:** Correspondence:* Annalisa Radeghieri, Department of molecular and Translational Medicine, University of Brescia, Viale Europa 11, 25123 Brescia, (Italy).

## Abstract

Antithrombin (AT) is a glycoprotein produced by the liver and a principal antagonist of active clotting proteases. A deficit in AT function leads to AT qualitative deficiency, challenging to diagnose. Here we report that active AT may travel physiosorbed on the surface of plasma extracellular vesicles (EVs), contributing to form the “EV-protein corona”. The corona is enriched in specific AT glycoforms, thus suggesting glycosylation to play a key role in AT partitioning between EVs and plasma. Differences in AT glycoform composition of the corona of EVs separated from plasma of healthy and AT qualitative deficiency-affected subjects were also noticed. This suggests deconstructing the plasma into its nanostructured components, as extracellular vesicles, could help to unravel pathophysiological mechanisms otherwise undiscovered.

## Introduction

Antithrombin (AT), a heparin cofactor and member of the serine protease inhibitor (serpin) gene family, is an important protease inhibitor that regulates the function of several serine proteases in the coagulation cascade^1^. AT physiologically inactivates thrombin (factor IIa) and factor Xa (FXa) and, to a lesser extent, factors IXa, XIa, XIIa, tissue plasminogen activator (tPA), urokinase, trypsin, plasmin, and kallikrein^2, 3^. The plasma concentration of AT is 112 to 150 mg/L, with a half-life of 2 to 3 days^4, 5^. The liver is the primary source of AT synthesis and post-translational glycosylation^6^. Mature AT has a molecular weight of 58 kDa and four potential N-glycosylation sites at asparagine (Asn) residues, occupied by a biantennary or triantennary structure of complex N-glycans bearing two terminal sialic acids. No other post-translational modifications are known to be present on the protein. Many isoforms of AT are documented, because of a microheterogeneity of attached carbohydrates. The 93-94% predominant group of isoforms is designated as α-AT while the 6-7% subsidiary group form as β-AT: they can be isolated thanks to different binding to glycosamminoglycans^7, 8^. Even if less abundant, β-AT has higher affinity for heparin and is more important in controlling thrombogenic events from tissue injury^9-12^. Clotting inactivation by AT is the consequence of the entrapment of the coagulation factors in a covalent and equimolar complex in which the active site of the proteases becomes inaccessible to its substrate^13^.

AT circulates in a form that has a low inhibitory activity. Under normal physiological circumstances, the anticoagulant effect of AT is accelerated at least a thousand times in the presence of heparin-like glycosaminoglycans, such as heparan sulphates, located on the vascular endothelium. Besides, the interaction of AT with the endothelium gives rise to an anti-inflammatory effect: it increases the production of the anti-inflammatory cytokine prostacyclin, which then mediates smooth muscle relaxation and vasodilatation and inhibits platelet aggregation^14^.

The evidence for the role AT plays in regulating blood coagulation is demonstrated by the correlation between inherited or acquired AT deficiencies and an increased risk of developing thrombotic disease. Inherited AT deficiency is divided into type I deficiency (T1), in which both the functional activity and levels of AT are proportionately reduced (quantitative deficiency), and type II deficiency (T2), in which normal antigen levels are found in association with low AT activity due to a dysfunctional protein (qualitative deficiency)^15^.

In the last years, together with many other proteins usually considered “soluble” factors^16^, AT has also been found associated with platelet-derived extracellular vesicles (EVs)^17^ extracted from plasma or serum, and cell culture media-extracted EVs^18, 19^. EVs are nanoparticles released by eukaryotic cells in the form of a lipid membrane that encloses proteins, nucleic acids, and metabolites. They are today considered the third way of cell communication - other than direct intercellular stimuli and paracrine secretion of active molecules^20^ - placing EV research as a key field within immunology, haematology, and cancer cell biology^21, 22^.

The involvement of circulating EVs in coagulation processes has been documented, but their role is still debated, since both coagulation factors and anticoagulant proteins have been found associated to blood EVs^23-27^.

This work presents the first detailed analysis of the association of AT isoforms to circulating plasma EVs that could possibly have repercussions on AT-related deficiencies studies and beyond.

## Materials and Methods

### Patients and blood sample collection

Ethical approval was obtained from the Ethical committee of Spedali Civili hospital (Brescia, Nr.NP4761). Patients and control subjects enrolled in the study provided written consent according to the Declaration of Helsinki. Tests were performed on plasma samples obtained from the Haemophilia Centre, Haemostasis and Thrombosis Unit at Spedali Civili (Brescia). Peripheral blood samples were collected from 3 patients diagnosed with T2 AT deficiency, and healthy subjects. EDTA was added and samples were immediately centrifuged at 4200 *g* for 10 minutes at 22°C. Samples were anonymized and stored at -80 °C until analysis. At the time of analyses, about 1 mL of plasma was thawed at room temperature (RT) and examined.

### EV separation protocols

#### Ultracentrifugation (UC) and sucrose gradient

Plasma EVs were isolated through UC and discontinuous sucrose gradient^28^. Briefly, 1 mL of plasma was processed with three subsequent steps of centrifugation at 4°C: 800 *g*, 16,000 *g* and 100,000 *g*, respectively for 30 minutes, 45 minutes and 2 hours. The supernatant was discarded, and the EV pellet was resuspended in 1 mL of buffer (250 mM sucrose 10 mM Tris-HCl, pH 7.4) and loaded at the top of a discontinuous sucrose gradient. The gradient was centrifuged at 100,000 *g* for 16 hours at 4°C (rotor MLS-50, Beckman Optima MAX). Twelve fractions of 400 μL were collected from the top of the gradient and pelleted by UC at 100,000 *g* for 2 hours at 4°C (Figure 1). Fractions were analyzed with SDS-PAGE and Western blot (WB).

**Figure 1.**
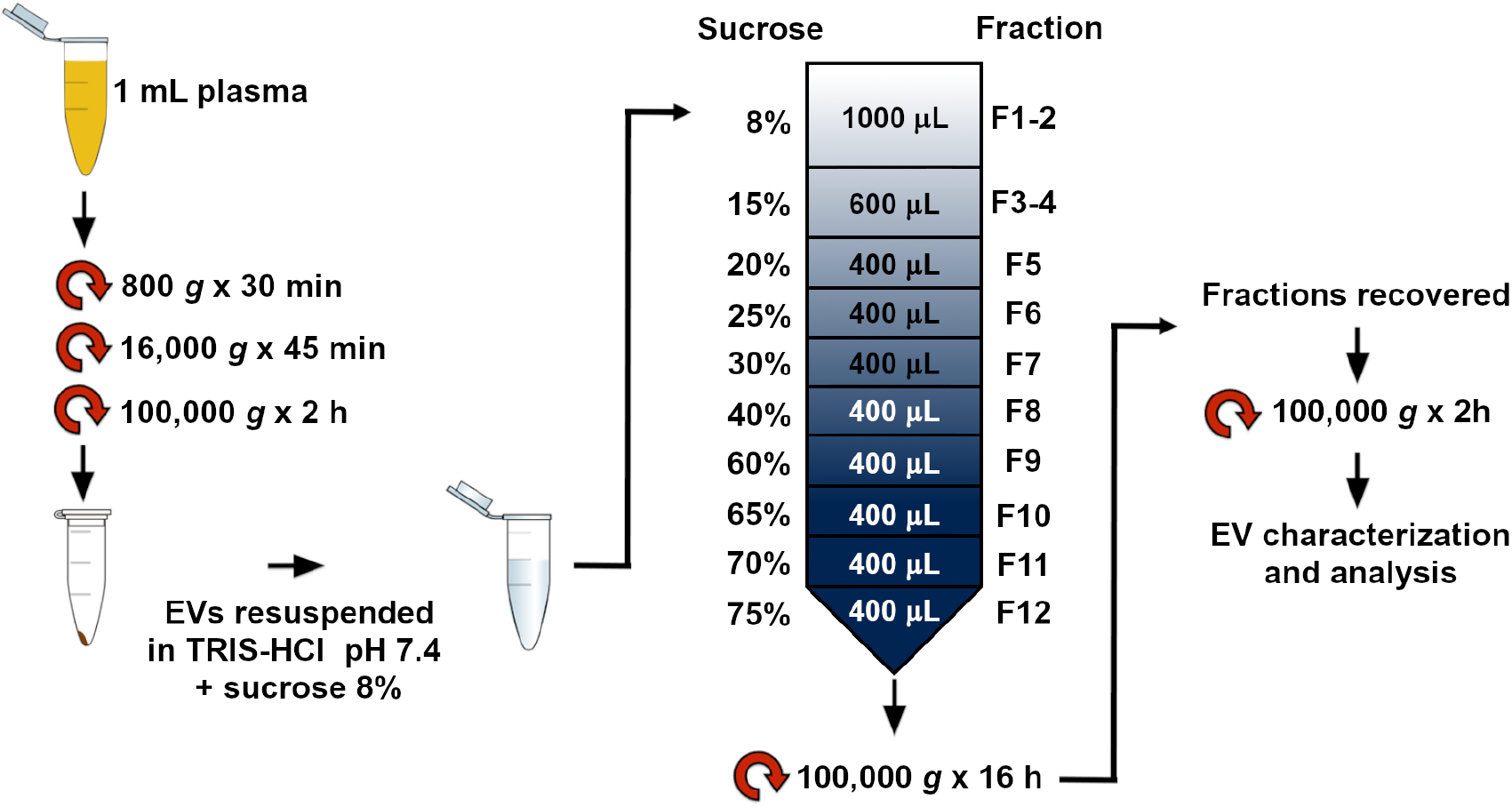
EV purification protocol. Key concept sketched. EVs were isolated from 1 mL of human plasma by differential centrifugation. The pellet was resuspended in Tris-HCl sucrose 8% and loaded on the top of a discontinuous sucrose gradient. Afterwards, twelve fractions were collected, pelleted by ultra-centrifugation and further analyzed.

#### Size exclusion chromatography (SEC)

Plasma EVs were also isolated through SEC, using IZON qEVsingle columns. We performed the separation following the producer datasheet on the EVs obtained by the UC steps. A hundred μL of the EV pellet in PBS were loaded on the top of SEC columns. Fractions of 200 μL were collected and fractions 6-11 were centrifuged at 100,000 *g* for 2 hours at 4°C. Pellets were resuspended and analyzed by SDS-PAGE and WB.

### EV characterization

#### Atomic Force Microscopy (AFM)

EV pellets were resuspended in 50 μL of PBS and diluted 1:10 v/v with deionized water. 5 to 10 μL of samples were then spotted onto freshly cleaved mica sheets (PELCO® Mica discs Grade V-1, thickness 0.15 mm, 10 mm diameter from Ted Pella, Inc). All mica substrates were dried at RT and analyzed using a Nanosurf NaioAFM equipped with Multi75AI-G probes (Budget sensors). Images were acquired in dynamic mode, scan size ranged from 1.5 to 15 μm and scan speed ranged from 0.8 to 1.5 seconds/line. AFM images were processed using Gwyddion ver. 2.58. The size of particles was extrapolated using built-in modules. Particle size distribution was then calculated on GraphPad PRISM ver. 6, by plotting particle size against relative abundance.

#### CONAN assay

EVs were resuspended in 100 μL of Milli-Q water to be checked for purity from protein contaminants and quantified using the COlorimetric NANoplasmonic (CONAN) assay we previously described^29^.

#### SDS-PAGE and WB

EV samples were resuspended in Laemmli buffer and boiled for 5 minutes, before being separated on a 10% polyacrylamide gel. Proteins were then transferred onto a polyvinylidene difluoride (PVDF) membrane (GE Health-care) for immunoblotting and blocked with 5% (w/v) fat-free dried milk in PBS 0.05% Tween-20 (PBST) for 1 hour at 37°C. Membranes were incubated overnight at 4°C with primary antibodies diluted in PBST 1% (w/v) fat-free dried milk. Membranes were washed 3 times in PBST and incubated with the HRP-conjugated secondary antibodies in PBST 1% (w/v) fat-free dried milk for 1 hour at RT. After three washes, chemiluminescence was acquired using Bio-Rad Clarity Western ECL on a G:Box Chemi XT Imaging system (Syngene)^30^. Primary antibodies used: sheep anti-ATIII (Hematologic Technologies INC, PAHAT-S), rabbit anti-Adam10 (Origene, AP05830PU-N), mouse anti-Alix (Santa Cruz, sc-53539), mouse anti-CD63 (Millipore, CBL553), mαCD81 (Santa Cruz, sc-7637), mouse anti-Tsg101 (Santa Cruz, sc-7964), rabbit anti-ApoI (Thermofisher, 701239), and mouse anti-GM130 (BD Bioscience, 610823). Secondary antibodies used: rabbit anti-mouse (Bethyl, A90-117P), donkey anti-sheep (Bethyl, A130-100P), and goat anti-rabbit (Bethyl, A120-101P). Primary antibodies were diluted 1:1000 except for sheep anti-ATIII (1:3000). Secondary antibodies were diluted 1:3000.

#### Dot blot assay and trypsin treatment

Dot blot was performed as previously described^31^. Briefly, EVs were resuspended in 100 μL of 100 mM Tris, 150 mM NaCl, 1 mM EDTA. 5 μL of EVs diluted 1:1, 1:2, and 1:5 (v/v) in buffer were spotted on a nitrocellulose membrane and allowed to dry at RT for 1 hour. Membranes were then blocked with 5% (w/v) fat-free dried milk in Tris-buffered saline (TBS) in the absence or presence of 0.1% (v/v) Tween-20 for 1 hour at RT, followed by the incubation overnight at 4°C with anti-CD63, anti-Ago2 (Origene, TA352430) and anti-ATIII antibodies, diluted 1:1000 in TBS or TBST 1% fat-free dried milk. After 3 washes with TBS or TBST, membranes were incubated with HRP-conjugated with proper secondary antibodies diluted in TBS or TBST 1% fat-free dried milk for 1 hour at RT. Blots were detected as described above. EV samples were also treated with 0.25% trypsin and incubated for 10 minutes at 37°C. Samples were then centrifuged at 100,000 *g* for 2 hours. The pellet and supernatant were spotted on the nitrocellulose membrane, as just described. The membranes were incubated with or without 0.1% (v/v) Tween-20.

### 2D Gel Electrophoresis (2D SDS-PAGE)

#### TCA-DOC/Acetone purification

To precipitate proteins and remove contaminants, plasma and EV samples were incubated at RT with 2% Na deoxycholate (DOC) (0.02% final concentration) and then with 10% trichloroacetic acid (TCA) for 15 minutes and 1 hour, respectively. Samples were centrifuged at 20,000 *g* for 10 minutes at 4°C, then 200 μL of ice-cold acetone was added, incubating the sample on ice for 15 minutes. The step was repeated, and the resulting pellet was dried by inversion. Samples were finally resuspended with 350 μL of Rehydration buffer (Urea 7 M, Thiourea 2 M, CHAPS 4% (w/v), Carrier Ampholyte 0.5% (v/v), DTT 40 mM, Bromophenol Blue 0.002%), for the following 2D-PAGE.

#### 2D-PAGE

For the electrofocusing (IEF) phase, immobilized pH gradient (IPG) strips of 18 cm having a pH range 4-7 (Ready Strips, Bio-Rad) were used and were rehydrated overnight at RT, adapting from^11^. IEF was then performed on a Multiphor II Electrophoresis System with ImmobilineDryStrip Kit (Amersham Biosciences, Ge Healthcare), in 4 consecutive steps: 1) 1 minute at 500 V, 1 mA, 5 W; 2) 1 hour at 500 V, 1 mA, 5 W; 3) 4 hours at 3500 V, 1 mA, 5 W; 4) 13.30 hours at 3500 V, 1 mA, 5 W. After IEF, strips were equilibrated at RT under gently mixing with two different solutions of Equilibration buffer (SDS 2% (w/v), Urea 36% (w/v), 50 mM Tris-HCl pH 8.4, Glycerol 30% (v/v)) with DTT 2% (w/v) and iodoacetamide 2.5% (w/v), for 12 and 5 minutes, respectively. Strips were then frozen. Subsequently, strips were cut and the 6 cm piece with a pH range of 5-6 was inserted on the top of 8% SDS-PAGE gel and sealed with hot agarose solution (0.5% agar and Bromophenol Blue in Running buffer 1X). Electrophoresis and subsequent Western blot were performed as described above.

### Antithrombin activity on EVs

Formation of thrombin-antithrombin complex (TAT complex) was evaluated by incubation of circa 1.5 ng of AT from plasma EV samples with 0.0025 U of thrombin at 37°C. Aliquots were withdrawn at different time intervals (30 minutes, 1 hour, 2 hours, 4 hours). The reaction was carried out with previous incubation of AT with 0.06 U of heparin for 30 minutes at 37°C. These samples were evaluated by SDS-PAGE as indicated previously.

### EV Track

All relevant data has been submitted to the EV-TRACK knowledgebase^32^. EV track ID: EV21008.

## Results

### 1. Antithrombin is associated to plasma small EVs

To separate and purify EVs from plasma, we performed differential (ultra)-centrifugations, followed by purification through a discontinuous sucrose gradient^28^.

Figure 2A shows a representative Western blot of the twelve fractions obtained from the gradient performed on a healthy plasma sample. To verify the presence of EVs, common EV markers as Adam10, Alix, Tsg101, CD63 and CD81^33^ were probed. The negative marker GM130, was analyzed to exclude cellular contamination in the samples, and ApoAI, the specific marker for HDL lipoproteins, was checked to verify the presence of HDLs in the sample.

**Figure 2.**
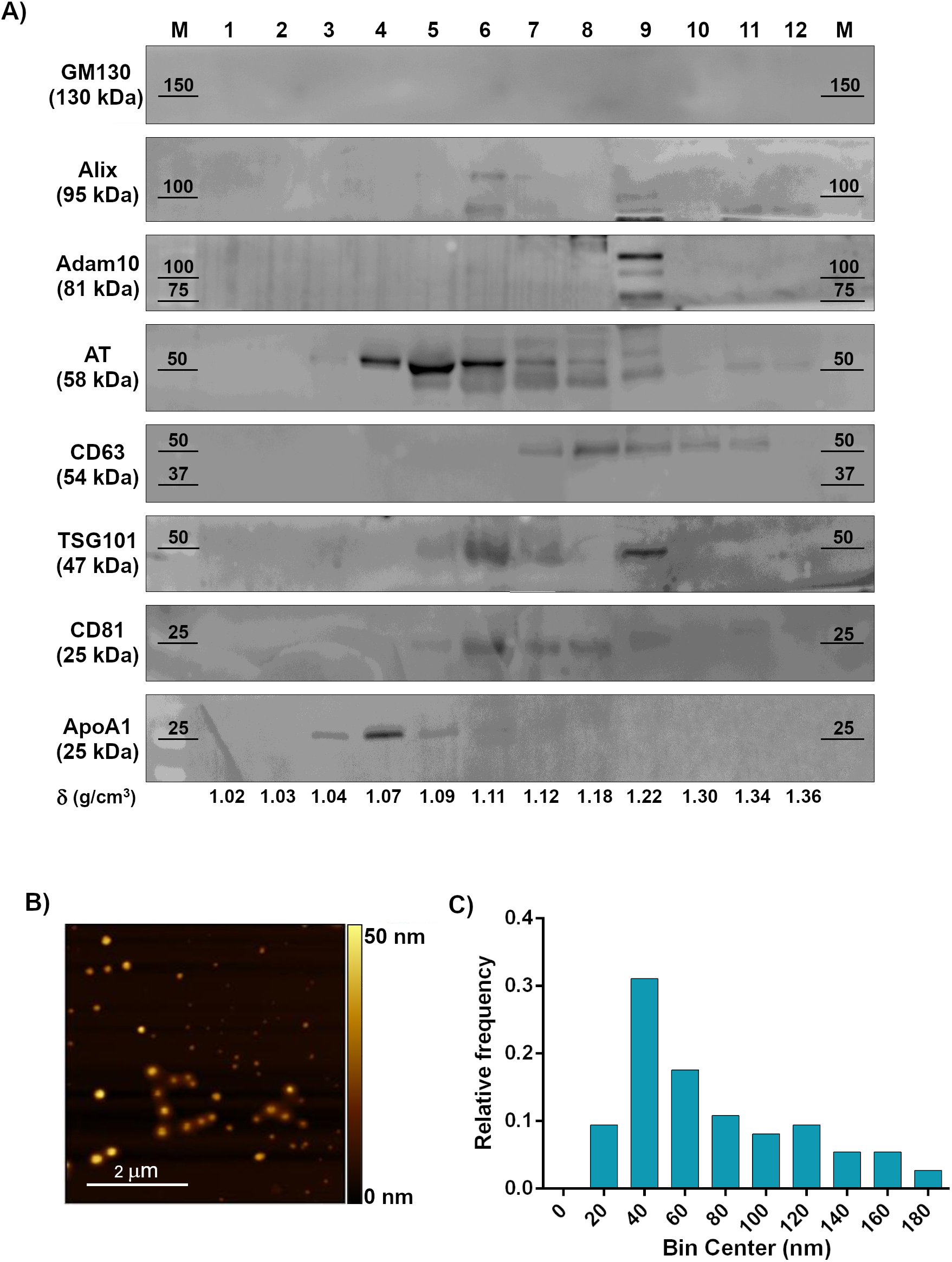
Comprehensive characterization of plasma-derived EVs. A) Western blot analysis after SDS-PAGE of sucrose gradient fractions in reducing conditions. The distribution of EV (Alix, Adam10, CD63, TSG101, CD81), Golgi (GM130, negative control) and lipoprotein (ApoA1, contaminant) markers is shown, together with AT. Fractions from 6 to 9 feature both the absence of contaminants and the co-localization of AT and EV markers. B) Representative AFM image of the particles in fractions 6-9, displaying intact and round shape objects, and a size comparable to the one of EV dried on a surface. C) Size distribution of the particles in fractions 6-9, imaged through AFM. Particle diameter was extrapolated from 200 objects.

Thanks to this purification protocol, we could observe an enrichment of EVs in fractions 6 to 9 (with density ranging from 1.11 and 1.22 g/cm^3^). GM130 is absent in all fractions, meaning that our samples do not contain any cellular contaminants. AT is enriched from fractions 4 to 9, overlapping the main EV markers. Recent studies demonstrated that the fraction with a density of 1.09 g/cm^3^ (corresponding in our gel to fraction 5) may contain protein macro-aggregates or ectosomes, vesicles with a cellular origin that incorporate the coagulation factors released by platelets and endothelial cells^34^.

The sucrose gradient allowed us to separate also HDLs, which are present in fractions 3 to 5 and are particularly abundant in fraction 4 (with a density of 1.07 g/cm^3^, comparable with the density of the most abundant HDL types^35^). Gradient fraction 4 shows the presence of ApoAI and AT: it might suggest that AT circulates in blood not only associated with EVs but also with HDLs. Afterwards, fractions 6-9 were pooled and used for the subsequent analysis steps, being the fractions most enriched in EVs and deprived in HDLs.

Pooled particles from fractions 6 to 9 were then imaged. AFM was employed to verify the presence of EVs in the selected fractions (6-9) and to visualize their morphology and size^36, 37^. Round-shaped objects with diameters ranging from 20 to 170 nm (Figure 2B) are visible in all the fractions tested. According to the size distribution analysis (Figure 2C) performed, most of the particles (∼ 95%) are comprised in the 30-180 nm range, which is compatible with the size of small EVs. Objects featuring a size < 30 nm are almost absent (< 5%), further confirming the content of smaller biogenic particles of the fractions (e.g. HDLs, often co-isolated with small EVs) is negligible.

To confirm that AT is present on EVs and that this finding is not an artefact of the separation method used^38^, we have also performed the separation of EVs by size exclusion chromatography (SEC), a more gentle and non-disruptive method. As seen in Figure S1, AT is present in EV fractions eluted from SEC together with EV markers Adam 10 and CD81. Taken together these results show that AT can circulate in blood on EVs and not only in a soluble free form^39^.

We next performed a semi-quantitative analysis to estimate the levels of AT attached to EVs. Figure 3A reports a representative Western blot of free AT in plasma at different dilutions (lanes 1 to 4) and EV-bound AT (“EVs” lane). Starting from the reported physiological plasmatic AT concentration, we stoichiometrically calculated the concentration of AT in each of the diluted plasma samples loaded on the gel (lanes 1 to 4) and related it with the corresponding band intensity. Finally, we plotted a calibration line (Figure 3B) and used it to estimate the amount of AT contained in the EV sample (Figure 3B, red star). From our calculations, it appears that the concentration of AT physiosorbed on EVs separated from 1 ml of plasma is between 3.7 and 4.0 pg, that corresponds to 0.0026% - 0.0032% of the total AT contained in 1 ml of human plasma.

**Figure 3.**
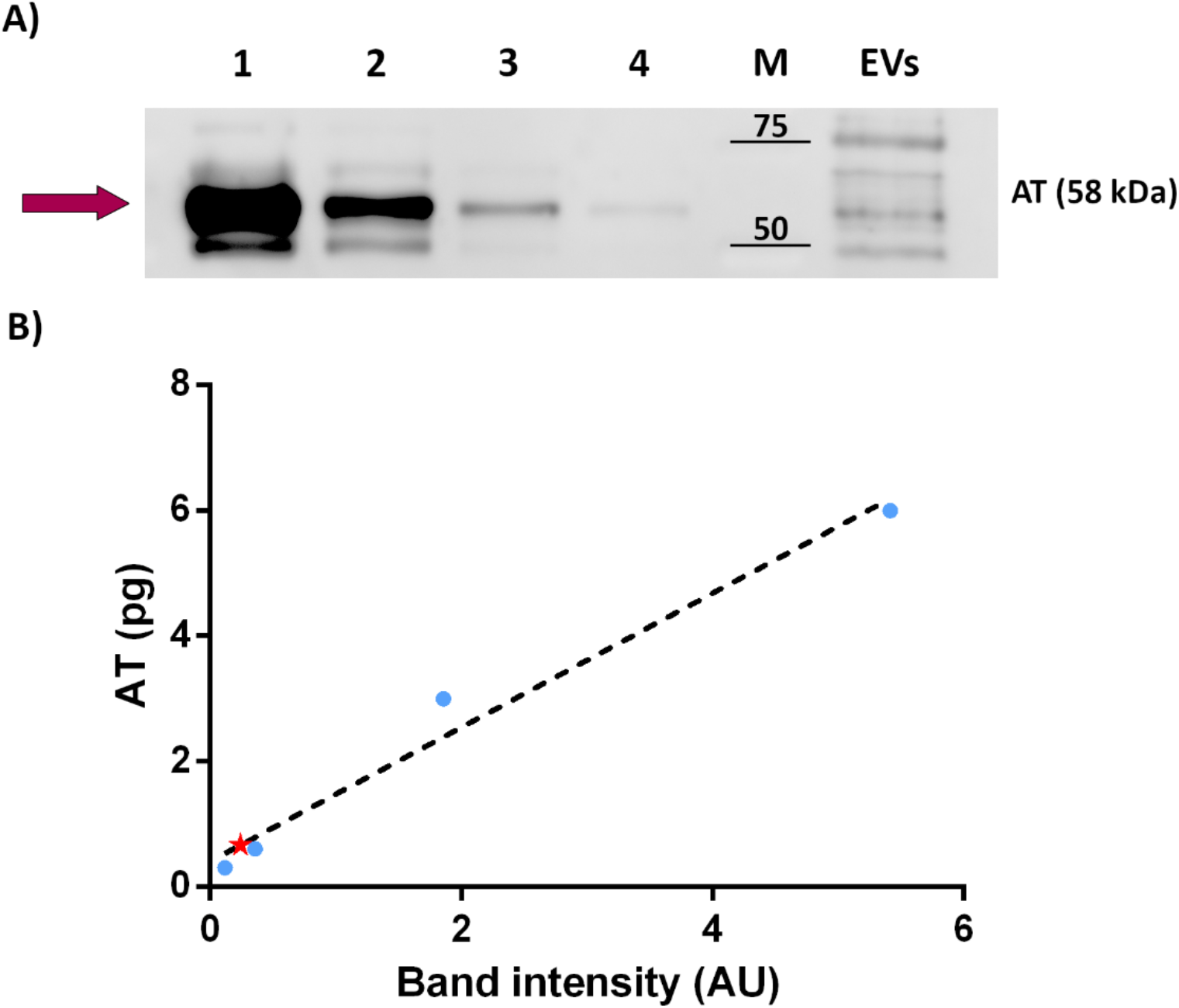
Semi-quantitative analysis of AT concentration on EVs. A) EV-bound AT was measured using Western blot. Diluted plasma samples were used as standards. Lanes from 1 to 4 contain a diluted amount of plasma. Total protein loaded: 2.56 μg in lane 1; 1.28 μg in lane 2; 0.25 μg in lane 3; 0.13 μg in lane 4. The “EVs” lane was loaded with 1,42 μg of proteins, corresponding to the EVs separated from 160 μl of plasma. AT band (58 kDa) is highlighted by the purple arrow. B) The calibration line created by plotting and linearly fitting (R^2^ = 0.9789) the amount of AT stoichiometrically calculated for plasma loaded in lanes 1-4 *versus* the corresponding band intensity measured through densitometric analysis. Blue dots: diluted plasma samples. Red star: EV sample.

### 2. AT is localized at the EV surface

AT is known in literature as a secreted protein circulating in the blood, although our evidence highlights it is also associated with EVs.

To characterize the nature of this binding, a dot blot analysis has been performed. The technique consists of spotting purified EVs onto a nitrocellulose membrane at different dilutions. Afterwards, a WB is performed. Under these experimental conditions, if the vesicular protein is exposed to the solvent, it should be detected in the presence or absence of a detergent (which disrupts the membranes and allows the inner proteins to spread in the solution). As opposite, if the protein is contained inside the vesicles, it should be detected only in the presence of the detergent^40, 41^. Argonaute2 (Ago2), being a protein present within the EVs, has been used as a negative control in this experiment. As shown in Figure 4A, only blots immunolabeled in the presence of the detergent revealed Ago2 signals when compared with the control (without detergent). On the contrary, the positive control CD63 was detected on both membranes treated or not with detergent. This EV marker is a transmembrane protein detected on the surface of EVs.

**Figure 4.**
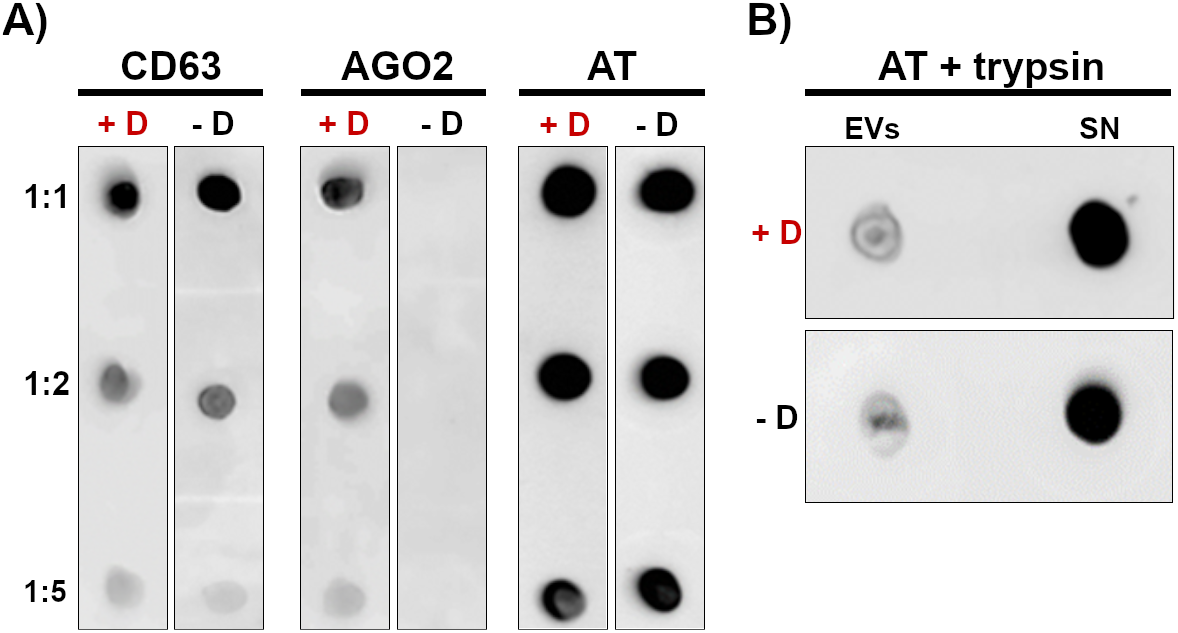
Analysis of AT localization onto EVs. EVs from plasma samples of healthy donors were spotted on nitrocellulose membrane and analyzed with dot-blot. All the tests were performed in presence (+D) and absence (-D) of 0.1% (v/v) Tween-20. A) Immunoblot vs. CD63 (EV membrane-associated protein), Ago2 (lumen protein), and AT in presence and absence of 0.1% (v/v) Tween-20. Membrane protein (CD63) signal is not affected by the detergent, while EV lumen protein (AGO2) is revealed only when the EV membrane is disrupted. The behavior of AT suggests the association with EV membrane rather than its encapsulation within EV lumen. B) Immunoblot vs. AT performed on plasma EVs and supernatant after trypsin treatment and in presence (+D) or absence (-D) of 0.1% (v/v) Tween-20. Supernatant enriches in AT after trypsin treatment, meaning AT is directly accessible to the protease action, and further suggesting its localization on the outer leaflet of EV membrane.

Results obtained with this experiment indicate that AT is localized in the EV membrane facing outwards since antibody anti-AT recognizes the epitope located outside of the vesicles, hence it cannot be an internal protein of EVs.

To verify whether AT could be detached from the EV membranes, EVs have been treated with an excess of trypsin (Figure 4B). Trypsin is a serine protease that hydrolyses proteins cleaving peptide chains mainly at the carboxyl side of the amino acid lysine or arginine. This treatment should detach and cleave the proteins linked or accessible in the vesicle membrane, releasing them in the extra-vesicular medium^42^. Purified EVs from the sucrose gradient fractions were resuspended in a specific buffer with the addition of trypsin. After incubation and separation by ultracentrifugation, the pellet and supernatant were then dotted on the nitrocellulose membrane by applying the protocol described. As shown in Figure 4B, AT is enriched in the supernatant, meaning that trypsin has digested the protein and released it in solution. This observation corroborates the findings described above, suggesting that AT localizes at EV surface.

### 3. AT on EVs retains anticoagulant activity

To determine if EV-AT retains anticoagulant activity, we evaluated the formation of a thrombin–antithrombin (TAT) complex in vitro as reported earlier^43^. As shown in Figure 5, which reports the formation of TAT complex at 37°C in presence of thrombin and heparin, we show an increase of TAT complex formation with time, with a corresponding consumption of free AT. It was interesting to notice that EVs separated from plasma already show the presence of TAT complex, presumably due to a scavenging property of EVs in blood.

**Figure 5:**
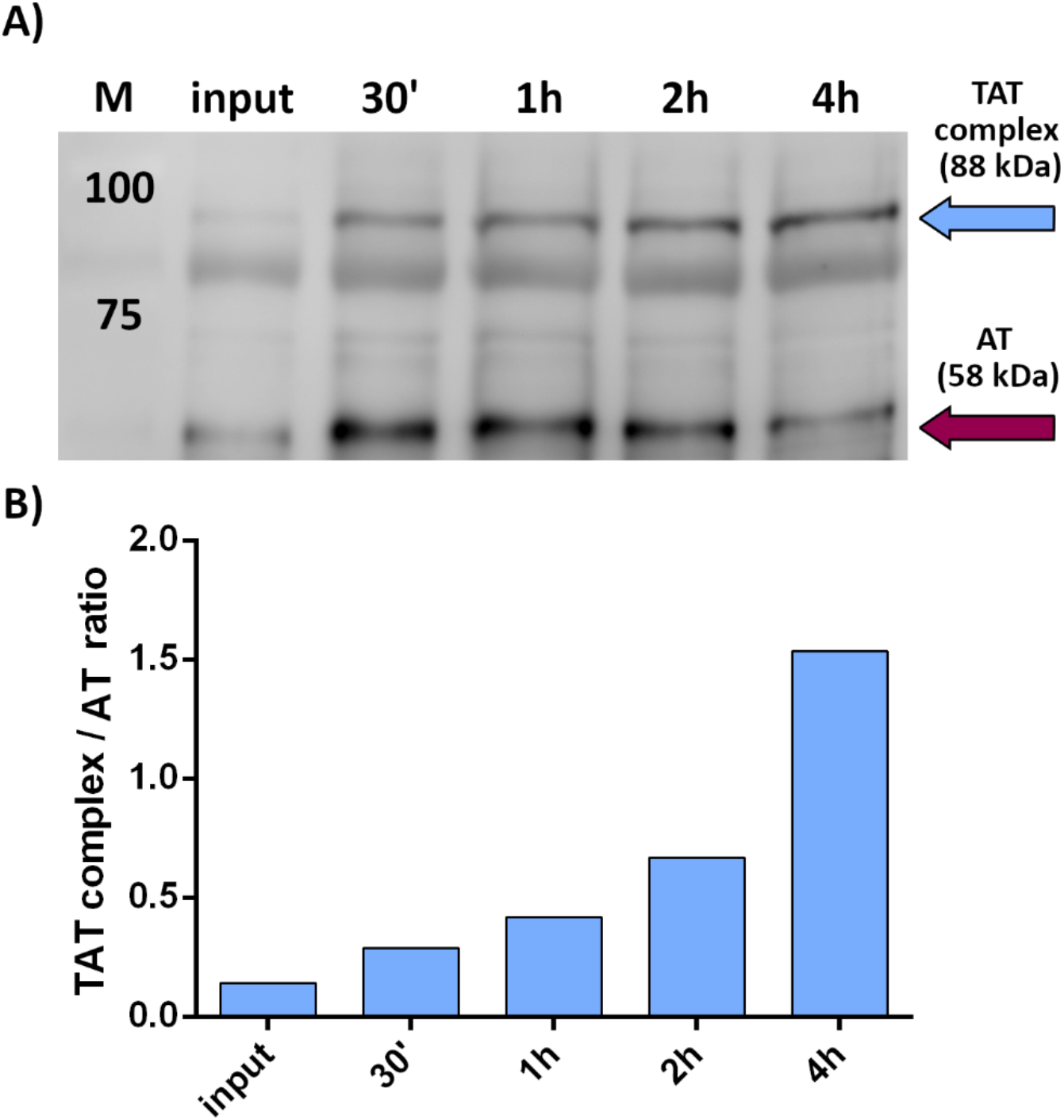
Determination of AT anticoagulant activity. EV-bound AT activity was assessed through the formation of the TAT complex. Fixed amounts of EVs (carrying ∼ 1.5 ng of AT) were challenged with 0.06 U of heparin and 0.0025 U of thrombin. A) TAT complex formation was measured at different timepoints through Western blot. Legend: cyan arrow = TAT complex band, purple arrow = AT band. B) Normalized TAT complex/AT ratio at different timepoints. For each timepoint, the TAT complex band intensity was normalized on the respective AT band intensity (posed equal to 1). The input corresponds to TAT complex/AT ratio at T = 0 minutes.

### 4. Specific AT glycoforms are enriched at the surface of EVs

As evidenced earlier, AT can be modified by N-glycosylation^12^. Since the partitioning of plasma proteins in blood seems to be influenced by glycosylation^44^, but no reported studies have compared EV glycosylation to the matched plasma, we verified if all the different glycoforms of AT circulating in plasma are also associated with EVs. To compare free soluble and EV-AT glycoforms we performed a 2D SDS-PAGE followed by WB on whole plasma (30 μg of proteins loaded on gel) and on isolated EV samples (gradient fractions 6-9), both derived from healthy subjects. Comparing the 2D electropherograms we could observe different patterns (Figure 6A). Soluble AT exhibits a particular pattern composed of many spots at different intensities, like what has been shown for purified soluble AT^11^. EV-AT pattern shows instead a different profile: the overall number of spots (glycoforms) is lower and there seems to be specificity in binding for certain glycoforms with respect to others. In particular, glycoforms between 5.05 and 5.15 in pI units (probably highly sialylated isoforms^11^) are almost absent in EVs. Furthermore, it is recognizable the presence of spots belonging to the group of β-AT isoforms (pI between 5.25 to 5.3). Overall, this pattern is not due to the total amount of loaded proteins but to a selective adsorption of those isoforms as evidenced in Figure 6C by the elaboration of densitometric profile of the AT spots of free and EV-AT, reporting relative abundances of AT glycoforms at different pIs intervals.

**Figure 6.**
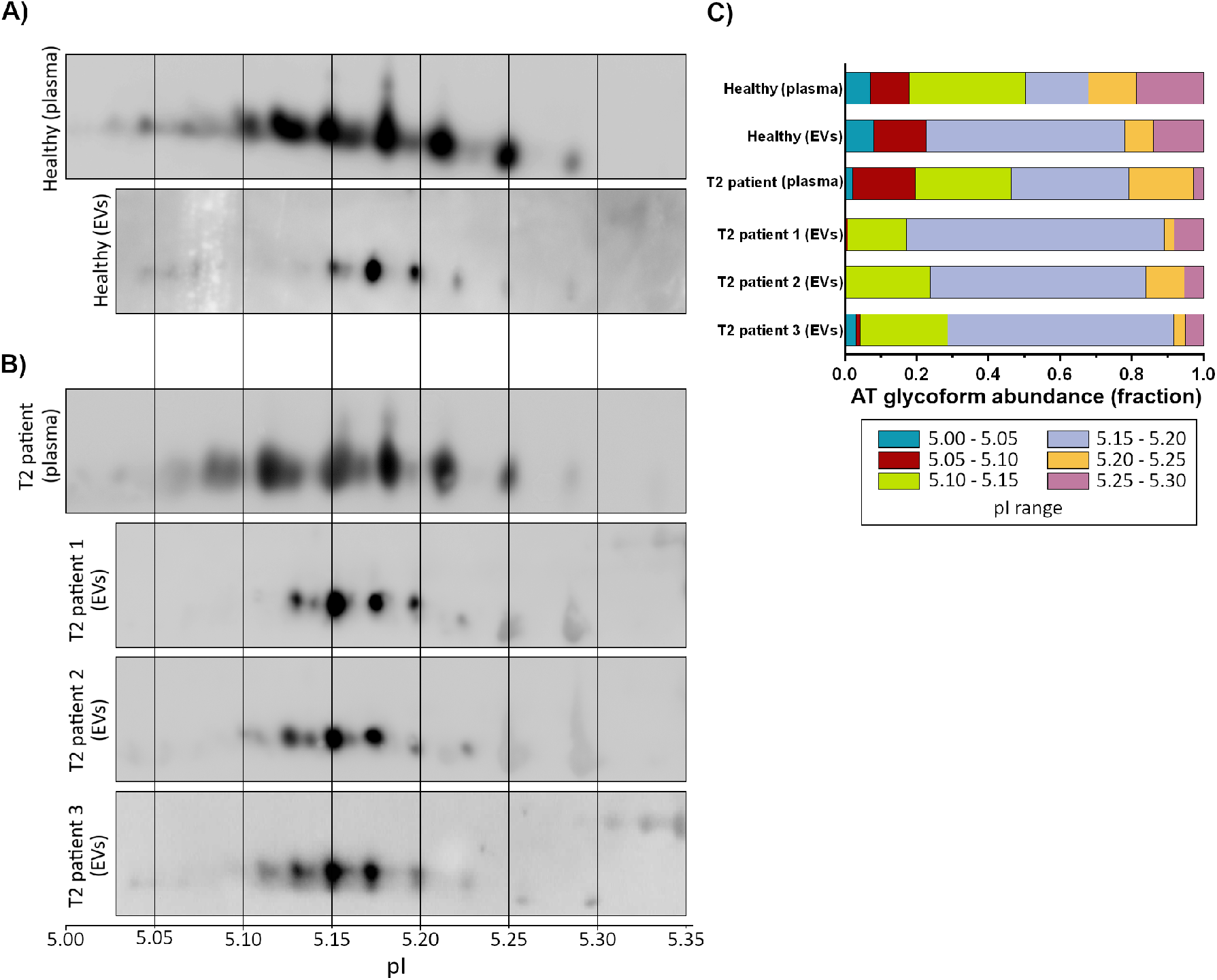
2D SDS-PAGE in reducing conditions on total plasma and isolated EV samples. A) Comparison of the 2D profile of soluble AT in total plasma (above) and EV-associated AT (below), both derived from samples of healthy subjects. pI is indicated. The number of spots is considerably different from plasma to EVs, suggesting a degree of “EV-specificity” of some AT glycoforms. B) 2D SDS-PAGE of whole plasma (one representative WB is reported) and EV-associated AT derived from T2 patients, showing a different migration pattern of the AT-EVs in respect to the control healthy sample. C) Densitometric analysis of the spots of soluble and EV-associated AT of healthy subjects and T2 patients. Differences in the relative abundance of AT glycoforms have been highlighted both in soluble and EV-associated AT of healthy subjects and in EV-associated AT of healthy subjects and T2 patients.

Hence, we suggest AT glycans are involved in the selective adsorption of AT to EVs.

### 5. EV associated AT shows a different 2D pattern between healthy and T2 patients

AT T2 deficiencies are caused by mutations within the primary sequence of AT, affecting either the reactive site or the heparin-binding domain or finally the mobility of the reactive loop after the binding with heparin^15^. Given that specific AT glycoforms selectively associate with EVs in healthy subjects, we verified if a mutation in AT coding sequence could lead to different AT adsorption patterns to EVs, possibly helping to discriminate between healthy and pathological samples and hopefully suggesting a possible molecular mechanism of AT deficiencies. We follow the hypothesis that mutations in the primary sequence of AT might lead to conformational changes in the structure of the enzyme, eventually altering the AT glyco-profile^43^, thereby influencing its adsorption to the EV surface^45, 46^.

We first performed a 2D SDS-PAGE of whole plasma from three T2 patients (all undergoing anticoagulant therapy), bearing the AT rare mutation R393C Northwick Park, which involves a reactive site defect. A representative WB of a patient plasma AT is shown in Figure 6B. By comparing the spot profile and the relative AT densitometric profile of patients with healthy subjects, we could not appreciate significant differences in spot distribution, apparently meaning that the mutation R393C does not influence the glycosilation profile of AT, nor the overall isoelectric point (as expected since the pI of Arg and Cys are very similar). We next performed a 2D SDS-PAGE of EV protein extracts derived from the same T2 affected patients. As shown in figure 6B, which reports representative WBs of the three EV samples obtained from patients probed with anti-AT, T2 samples present a higher number of EV-AT spots bearing a more basic pI, in respect to controls (healthy EVs, Figure 6A). In particular, the pI interval between 5.10 and 5.15 shows 2 to 4 spots that are not present in EVs from controls. The differences are also well evidenced in the elaboration of the densitometric profile shown in Figure 6C.

## Discussion

In the past few years, it has become clearer and clearer that EV surface molecules are of critical functional significance^16^ since they allow to establish connections with cells^47^ and with other biogenic nanoparticles in biological fluids^48^. EV surface molecules comprise integral, peripheral, and lipid-anchored membrane proteins, but also an extravesicular cargo of proteins adsorbed to EVs, at least partly recruited in body fluids after vesicle shedding.

We might assume that only few of the proteins on EV surface are recruited during vesicle biogenesis and/or release by the producing cell. Indeed, a s demonstrated for synthetic nanoparticles^49^, EVs can recruit on their surface numerous proteins which are present in circulation. The nature of such “protein corona” has recently started to be investigated as for other biogenic nanoparticles^16, 40, 50, 51^. Such investigation promises to disclose new properties of EVs since EV protein corona might be involved in EV-mediated cellular communication or can even provide EVs with new regulatory functions.

In this pilot study we provide strong evidence that active AT is part of the protein corona of EVs with a composition modulated by glycosylation. First, we have evidenced AT is effectively present on EVs, by applying different separation techniques, which allowed us to confirm the presence of the protein on-board. Since by bioinformatic predictions and literature data AT is not predicted to be a membrane-anchored protein and it is normally shed into the circulation by hepatocytes by transport along the exocytotic pathway^52^, we verified its association to EVs by dot blot analysis. This technique confirmed that AT shows an exofacial topology and it is indeed detachable by treatment with trypsin^42^. We next verified that physiosorbed EV-AT still retains activity evaluating the formation of the TAT complex *in vitro*. Not surprisingly, but never shown before, we noticed that EVs already carry the TAT complex on their surface, suggesting a role for EVs of “scavenger” bio-nanoparticles. Moreover, we wanted to quantify how much AT travels physiosorbed to EVs, and we have shown that around 0.01% of the whole plasma AT is effectively and actively travelling on EVs, a not negligible contribution.

As mentioned earlier, AT is a glycoprotein, and its glycosylation is important for its function as anticoagulant agent^45, 53^. Glycosylation is the only post translational modification(PTM) of the protein and no other modification has been reported in literature^11^. Glycosylation has also been evidenced to have a role in recruiting proteins to HDLs^44^, hence we verified if this PTM could also play a role in the association process of AT to EVs.

By performing 2D SDS-PAGE on healthy samples, which allows discriminating the different glycoforms of AT, we have evidenced for the first time that not all the free soluble glycoforms are equally adsorbed to EV surface, and some are more favoured for EV recruitment, indicating glycan specificity in recruiting AT onto EVs. By comparing the spot profile of EV-AT isoforms with previous published data^11^ we suggest that both α-AT isoforms and β-AT isoforms are present on EVs, even if at different ratio in respect to plasma. This is the first observation that a protein is specifically attached to the EV surface based on its glycosylation, although others have shown that glycosylation is important for EV biodistribution^54^ and functions^55^.

We believe that studying the selectivity of adsorption of AT glycoforms to EVs could be of importance to unravel hidden roles of AT in the coagulation process^56^. EVs could indeed offer a surface to accelerate the anticoagulant effect of AT in circulation, as it happens for vascular endothelium heparan sulfates^14^, or bring a local anti-inflammatory effect.

Notably, AT can be defined as a very “sticky” protein since it has also been found attached to the surface of HDLs^57, 58^, conferring these lipoproteins a direct role in coagulation that has not yet been investigated, and as protein corona component of PEG-liposomes after incubation with fetal bovine serum under dynamic and static conditions^59^. Since AT glycosylation has not been considered in these cited literature results, we might assume that a signature of specific AT glycoforms can be found on the protein corona of HDLs and synthetic liposomes, bringing along diagnostic and therapeutic implications.

We further verified whole plasma and EV-AT 2D electrophoretic profiles in T2 patients bearing the R393C mutation, which alters the formation of a hydrogen bond in the structure of the serpin, thus leading to higher K^d^ for heparin. No other mutations have been described for those patients. As evidenced by our analysis, whole plasma AT from T2 patients has a very similar spot profile compared to healthy controls, while EV-AT electropherogram shows that specific isoforms are adsorbed onto EVs in respect to controls, with a prevalence of more acidic glycoforms adsorbed to EVs in T2 patients (Figure 6B). All together this data let us hypothesize that the 2D profile of AT from whole plasma in T2 patients might be only apparently similar to controls, with isoforms that exhibit the same pI value but most probably have different glycan structures. This hypothesis surely needs confirmation by performing glycoproteomics on EV-AT and further structural studies. However, from our data, we can suggest that analysis of EV-EV[1], but in general analysis of EV protein corona, brings about another “dimension”, i.e. EV adsorption capacity, in our case related to glycosylation, which can better help to stratify protein structural alterations. As a general observation, we can conclude that the study of protein binding specificity on EV corona is a challenging but promising approach that could help to discover novel physiological roles of proteins and their involvement in different pathologies.

In a similar fashion, HDL glycoprotein composition, including specificity in glycosylation, can help to differentiate pathologies and correlate with HDL functions^61^.

Even if this study is still at its infancy, and new patients need to be screened, a future possible fall-out of these findings could be the setting up of new diagnostic tests based on EVs to identify AT deficiency subtypes, that could improve AT deficiencies diagnosis and management. Indeed, current methods to measure levels of functional AT make use of synthetic substrate technology^62^ bearing many limitations that also hampers the prognosis, since different subtypes may have a lower risk of thrombosis.

Our work wants to open a new perspective in the EV field, introducing the involvement of PTMs in the formation of EV protein corona, with promising perspectives for studies on pathologies related to post-translational defects (e.g. congenital disorders of glycosylations).

## Abbreviations

EVs: extracellular vesicles
AT: antithrombin
UC: ultracentrifugation
AFM: Atomic force microscopy

## Acknowledgments

This research was supported the University of Brescia through “Fondo per la ricerca ex 60%”, by MIUR through PRIN 2017E3A2NR_004 project to A.R and P.B. and by the National Interuniversity Consortium of Materials Science and Technology (INSTM) through internal funds.

## Authorship Contributions

Conceptualization: Annalisa Radeghieri, Doris Ricotta. Investigation: Annalisa Radeghieri, Silvia Alacqua, Francesca Todaro, Vanessa Previcini, Andrea Zendrini. Resources: Giuliana Martini. Writing – Original Draft: Annalisa Radeghieri, Silvia Alacqua, Andrea Zendrini. Writing-Review & Editing: All authors. Visualization: Andrea Zendrini. Supervision: Annalisa Radeghieri, Paolo Bergese. Project administration: Annalisa Radeghieri. Funding acquisition: Annalisa Radeghieri, Paolo Bergese.

## Disclosure of conflicts of interest

No potential conflict of interest was reported by the authors.

We thank Lucia Paolini, Giuseppe Pomarico and Valerio De Stefano for helpful discussions.

## Supplementary figures

**Figure S1.**
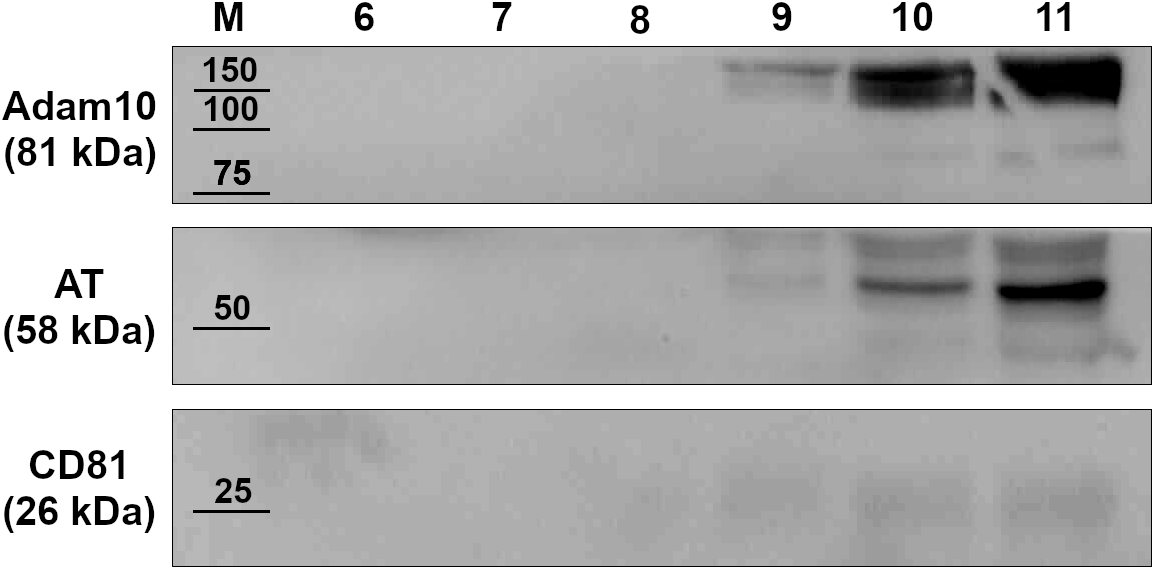
Biochemical characterization of plasma EVs obtained from size exclusion chromatography. The Western blot analysis performed after SDS-PAGE under reducing conditions of EV isolated with SEC confirmed the co-localization of the EV markers Adam 10 and CD81 with AT.

## Notes

### Competing Interest Statement

The authors have declared no competing interest.

### Summary of Updates

A functional test for AT has been introduced in the manuscript, which now shows that Antithrombin physiosorbed on EVs is active. A semi-quantitative analysis has also been attempted.

